# BIOLOGICAL DIAGNOSTICS OF THE ECOLOGICAL STATE OF URBAN ECOSYSTEMS

**DOI:** 10.1101/2022.11.21.517293

**Authors:** L.V. Galaktionova, N.A. Terekhova, N.G. Vedeneeva

## Abstract

The processes of urbanization and the development of industrial production entail active environmental pollution with pollutants of various nature. The existing analytical methods for their determination in natural media make it possible to reveal their concentration. But in conditions of complex soil pollution, this approach does not give a complete picture of their total impact on living organisms. In the course of the study, an assessment was made of the ecological state of soils in large administrative and industrial centers of the Southern Urals (the cities of Orenburg and Mednogorsk) in terms of their phytotoxicity and enzymatic activity. On the territory of the objects of study, soil samples were taken from a layer of 0-20 cm in various functional zones. Phytotoxicity was assessed using a wheat test culture, and indicators of urease and catalase activity were used to assess biological activity. The calculation of the integral indicator of the ecological state on their basis showed the development of processes of degradation of the soil cover of cities under the influence of urbanization.

## INTRODUCTION

The soil, as one of the geospheres, occupies a dominant role in the life of mankind. The close relationship between biotic and abiotic components, together with the growing pressure of the technogenic press, causes the need for biological diagnostics of soils. The biological components of the ecosystem contain information about the processes occurring in the soil environment, and are also capable of characterizing its danger with the help of their reaction to an external stimulus. Unlike analytical methods of pollution control, biological methods clearly show the effect of a complex of pollutants on the body, and, consequently, on the ecosystem as a whole [1].

In urban conditions, anthropogenic factors increasingly prevail over natural ones, so studying only the qualitative and quantitative composition of pollutants is not enough. Active urbanization leads to an increase in the degree of environmental risk for all components of natural ecosystems: air, vegetation, soil, water and ground [6]. Soil pollution has the strongest effect on soil formation as a result of the ingress of pollutants of non-soil origin into it [7].

Studies conducted by a number of authors indicate that the main carrier of heavy metals (HM) in urban ecosystems is the atmosphere, and the main sources are enterprises for the production and processing of non-ferrous metals, industrial waste and vehicles.

The specific effect of heavy metals on soil microorganisms is manifested in the inhibition of a number of biochemical processes, metabolic disorders. In general, the structure of the soil microbial community is restructured and the intensity of a number of physiological and biochemical processes (nitrogen fixation, nitrification, humification, and others) is reduced [5].

The ability of heavy metals to complex formation determines their ability to reduce the activity of soil enzymes with increasing concentration in the soil solution. Despite the different susceptibility of enzymes to the action of metals, their activity is a universal and informative marker of the development of techno genesis.

The territory of the Orenburg region, which is rich in natural resources and a developed agro-industrial complex, belongs to the zone of active technogenic impact. The list of pollutants varies widely, but heavy metals have remained a priority for 40 years. Therefore, studies on the assessment of the ecological state of the urban environment by methods of biological diagnostics are very relevant for the region [2].

## MATERIALS AND METHODS

The studies were conducted on the territory of large industrial and administrative centers of the Orenburg region – the cities of Orenburg and Mednogorsk in 2018-2022 (Figure 1).

**Figure 1.**
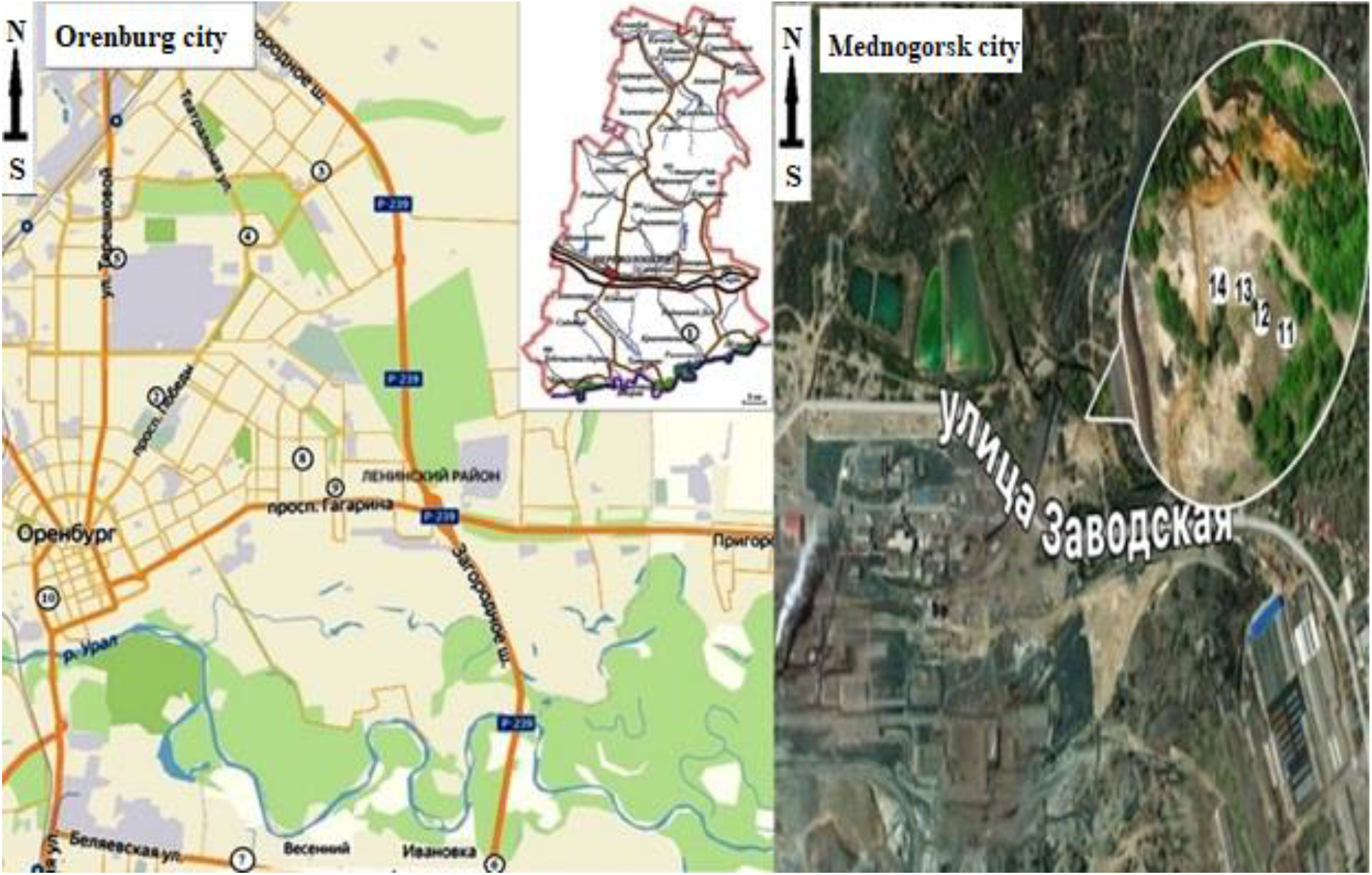
Soil sampling points Note: 1 - Background; 2 - Park them. Yu. A. Gagarin; 3 - gas station on Belyaevsky highway; 4 - Belyaevskoe highway; 5 - Lugovaya Street; 6 - Garden them. M. V. Frunze; 7 - Rosneft gas station; 8 - gas station Bashneft; 9 - Victory Avenue; 10 - New Street; 11 - Distance from LLC “MMSK” 50 m; 12 - Distance from LLC “MMSK” 100 m; 13 - Distance from LLC “MMSK” 150 m; 14 - Distance from LLC “MMSK” 250 m.

The soil cover of the study areas is represented by zonal soil types - southern chernozem with varying degrees of anthropogenic load. The background site is located 1–2 km northeast of the village of Krasnopole (Perevolotsky district, Orenburg region). Sampling was carried out using the “envelope” method at each registration site from a layer of 0–20 cm.

Phytotoxicity was assessed based on the assessment of germination and seed germination energy of *Triticum aestivum L*. according to GOST R ISO 22030-2009. The determination of catalase activity was carried out using the gasometrical method. The urease activity of soils was determined calorimetrically according to the method of T.A. Shcherbakova (1983) [3].

Statistical analysis of the acquired information was carried out using conventional methods and a package of practical projects MS Excel for Windows, Statistica V8 (StatSoft Inc., USA).

## RESULTS AND DISCUSSION

Data on the content of individual polluting components in the soil do not provide a complete picture of their total impact on the biotic component of ecosystems. Plant test objects are very sensitive to changes in environmental factors and make it possible to evaluate the integral effect of the toxic effect of soil components on higher plants.

High concentrations of HMs in the soil can have a phytotoxic effect. Plants of various systematic groups are used as test objects for biological diagnostics.

According to the data obtained, the soils of the city of Mednogorsk, located in the area of operation of LLC «MEDNOGORSK COPPER-SULFUR COMBINE», and the soils of Orenburg, located in close proximity to gas stations (Figure 2), have the highest phytotoxicity.

**Figure 2.**
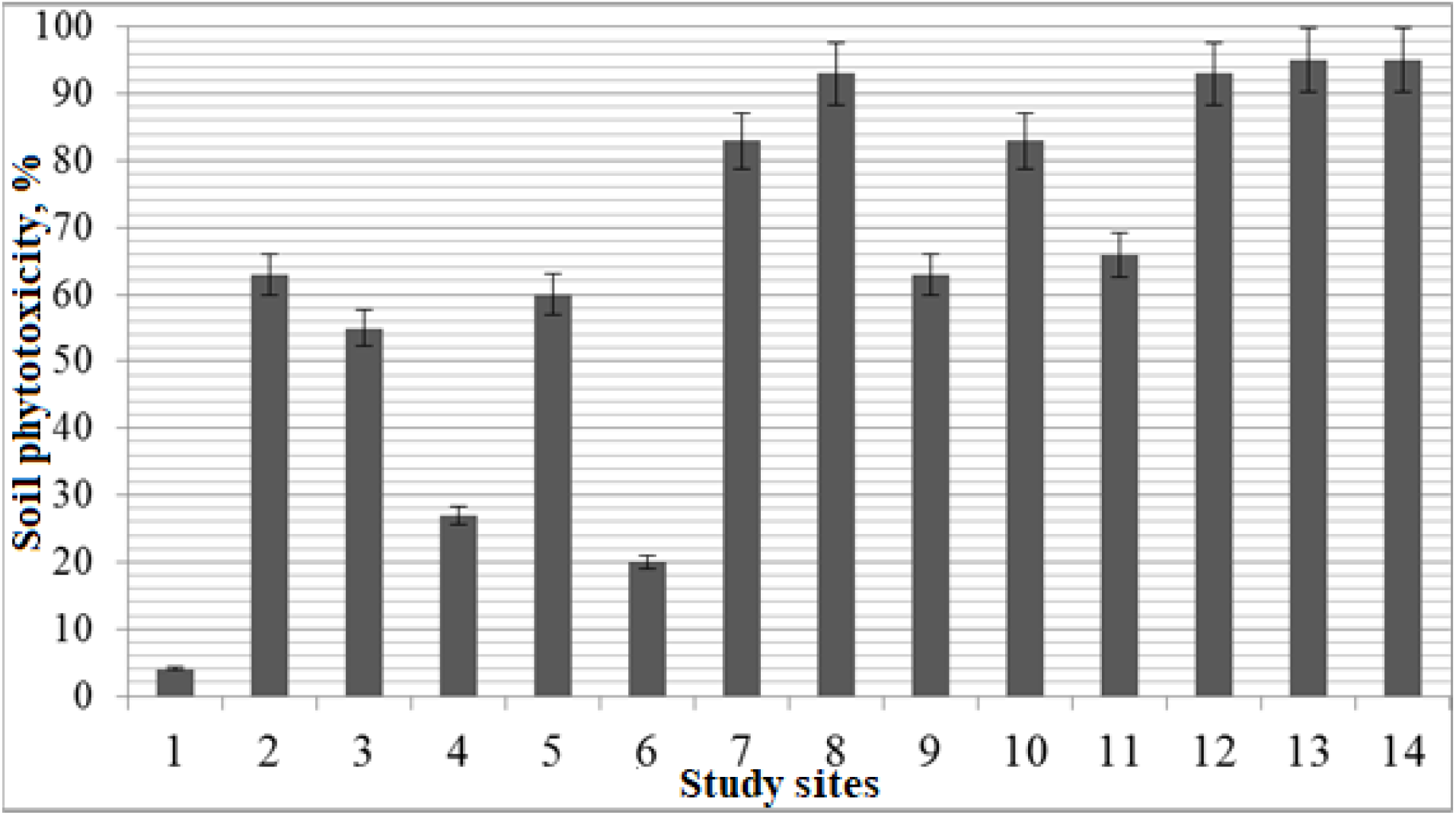
Phytotoxicity of soils in the cities of Orenburg and Mednogorsk.

The soil cover of the area located along the Belyaevsky highway in Orenburg (27,2%) is assessed as slightly toxic according to the scale proposed by Popova E.I. [10]. The soils of other sites were characterized as highly toxic to the test culture (Novaya Street, Rosneft and Bashneft gas stations) and hypertoxic (sites adjacent to MMSK LLC at a distance of 100 to 250 m). The rest were characterized by soils with an average level of toxic effects on *Triticum aestivum*.

Soil enzymatic activity is controlled by a number of different factors, one of which is heavy metals [4].

Urease is one of the most studied indicators and plays an important role in the conversion of soil nitrogen. The presence of urease in bacteria enables them to use urea as a source of ammonium, since it catalyzes its hydrolysis. Each soil cover has its own stable level of urease activity, which is determined by the ability of soil colloids, mainly organic ones, to exhibit protective properties by immobilizing enzymes [8].

The activity of urease in the study areas ranged from 14,0 to 47,9 mg N-NH_4_ per 1 g of soil in 4 hours. The average value for the city of Orenburg was 37,6 mg N-NH_4_ per 1 g of soil for 4 hours, and for the city of Mednogorsk – 20,65 mg N-NH_4_ per 1 g of soil for 4 hours, which is associated with more severe pollution of the soil cover cities operating enterprises of the metalworking industry. The activity of the enzyme in the background plot was 39,6 mg N-NH_4_ per 1 g of soil for 4 hours, which corresponds to the scale of soil enzymes by D.G. Zvyagintsev (Figure 3).

**Figure 3.**
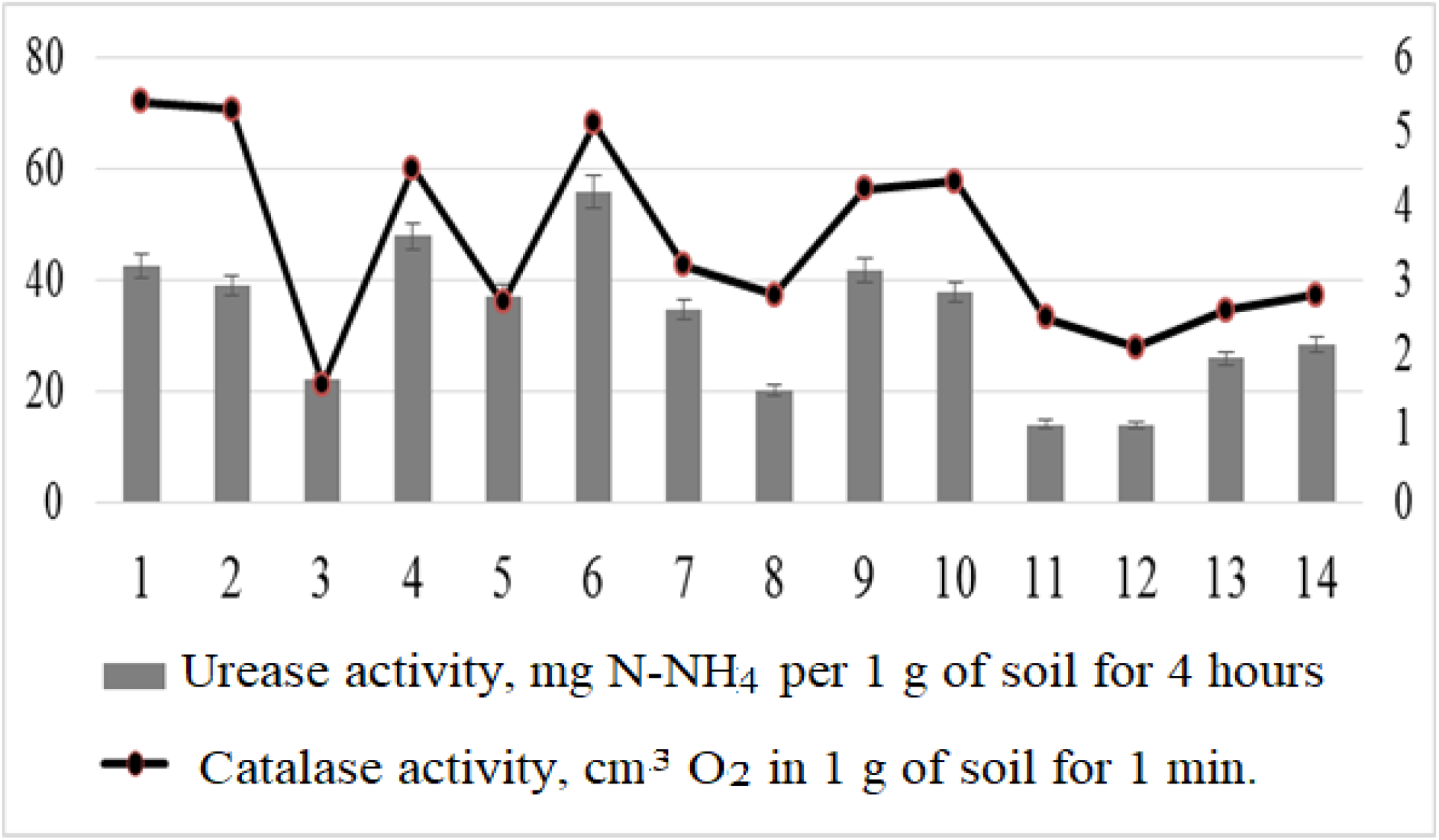
The activity of soil enzymes in the soils of the cities of Orenburg and Mednogorsk.

Soils poor in the degree of enrichment with urease are plots located near the Bashneft gas station and adjacent to MMSK LLC.

Enzymes related to oxidoreductases take an important part in redox reactions in the synthesis of humic substances. Catalase is one of the enzymes that is very responsive to changes in the intensity of anthropogenic load. Another factor affecting the activity of this enzyme is the presence of vegetation cover and its structure. The presence of a powerful root system capable of penetrating the soil layer significantly increased the activity of catalase in the soil. There are a number of regularities that significantly affect the change in the activity of the enzyme, so a direct relationship was found with a change in temperature, and the opposite was found with fluctuations in soil moisture. However, when moving down the profile, no significant changes in enzyme activity are observed [9].

The research results showed that catalase activity varies from 0,9 to 5,3 cm^3^ O_2_ per 1 g of soil per 1 minute. The average value for the city of Orenburg is 3,51 cm^3^ O_2_ per 1 g of soil per minute, for Mednogorsk – 2,9 cm^3^ O_2_ per 1 g of soil per minute (Figure 4). The background plot was characterized by high biological activity of the soil, since there are no industrial enterprises there and indirect anthropogenic impact is insignificant. The activity of the enzyme in the background area was 5,4 cm^3^ O_2_ per 1 g of soil in one minute, which corresponds to the average enrichment of soils according to the scale of D.G. Zvyagintsev. The soils of the plots were characterized by low enrichment with the enzyme: gas station on Belyaevsky highway, st. Lugovaya, Orenburg, and sites located in the coverage area of OOO MMSK, Mednogorsk.

On the basis of the data obtained by us, an integral indicator of the biological state of soils (IPBSP) was calculated, which made it possible to assess the ecological state of the soil cover of the study areas.

The results obtained indicate that 61,5% of the study sites are classified as highly degraded soils, and the rest are moderately degraded.

The assessment of the content of heavy metals in the soils of the study areas showed an excess of the MPC in the areas located in the zone of influence of the gas station and LLC “MMSK” in Mednogorsk. The calculation of the Zc indicator showed that in the territory of Orenburg, the sites dedicated to the gas station and along the street Lugovoi, are characterized by a dangerous category of soil pollution. And the sites located in the city of Mednogorsk, dangerous and extremely dangerous categories of pollution.

Statistical processing of the obtained results made it possible to reveal a significant correlation dependence of the activity indicators of soil enzymes with the content of copper, zinc and lead, as well as with the integral indicator of soil pollution Zc. Thus, the activity of soil catalase significantly correlated with the concentration of copper (r = – 0,49, p ≤ 0,05) and lead (r = – 0,77, at p ≤ 0,05) in soils, and urease activity only with the content Pb (r = – 0,51, p ≤ 0,05). The calculated correlation coefficient between Zc and catalase activity was -0,75 (p≤0,05), and urease -0.71 (≤0,05).

## CONCLUSION

Thus, at all study sites, regardless of their territorial location, there is a decrease in the integral indicator of the biological state of soils, which indicates violations of ecosystems under the influence of the urban environment. And the calculation of integral indicators of the biological state and pollution (Zc) of soils with heavy metals showed similar results, which allows us to conclude that the methods of biological diagnostics of the ecological state of soils in urban areas are highly informative.

## ACKNOWLEDGMENTS

The research was funded by the Ministry of Science and Higher Education in accordance with the state assignment on science for Ural State Mining University №075-03-2022-401 dated 12.01.2022 ‘Development and environmental and economic substantiation of the technology for reclamation of land disturbed by the mining and metallurgical complex based on reclamation materials and fertilizers of a new type’. We obtain the scientific results with the staff of Center for the collective use by using the equipment of the Center for the collective use of scientific equipment of the Federal Scientific Center of biological systems and agricultural technologies of RAS as well (No Ross RU.0001.21 PF59, the Unified Russian Register of Centers for Collective Use - http://www.ckp-rf.ru/ckp/77384).

